# Modular co-option of cardiopharyngeal genes during non-embryonic myogenesis

**DOI:** 10.1101/443747

**Authors:** Maria Mandela Prünster, Lorenzo Ricci, Federico Brown, Stefano Tiozzo

**Affiliations:** Sorbonne Université, CNRS, Laboratoire de Biologie du Développement de Villefranche-sur-mer (LBDV), 06230 Villefranche sur Mer, France.; Harvard University, Department of Organismic & Evolutionary Biology, 52 Oxford Street Cambridge, MA 02138 US; Departamento de Zoologia - Instituto Biociências, Universidade de São Paulo, São Paulo, SP, Brazil, CEP 05508-090; Centro de Biologia Marinha (CEBIMar), Universidade de São Paulo, São Sebastião, SP, Brazil, CEP 11612-109

**Keywords:** Blastogenesis, ascidians, *Botryllus schlosseri*, Budding, evo-devo, regeneration

## Abstract

**Background:** In chordates cardiac and body muscles arise from different embryonic origins. Myogenesis can in addition be triggered in adult organisms, during asexual development or regeneration. In the non-vertebrate ascidians, muscles originate from embryonic precursors regulated by a conserved set of genes that orchestrate cell behavior and dynamics during development. In colonial ascidians, besides embryogenesis and metamorphosis, an adult can propagate asexually via blastogenesis, skipping embryo and larval stages, and form anew the adult body, including the complete body musculature.

**Results:** To investigate the cellular origin and mechanisms that trigger non-embryonic myogenesis, we followed the expression of ascidian myogenic genes during *Botryllus schlosseri* blastogenesis, and reconstructed the dynamics of muscle precursors. Based on the expression dynamics of *Tbx1/10, Ebf, Mrf, Myh3* for body wall and of *FoxF, Tbx1/10, Nk4, Myh2* for heart development we show that the embryonic factors regulating myogenesis are only partially co-opted in blastogenesis, and propose that the cellular precursors contributing to heart or body muscles have different origins.

**Conclusions:** Regardless of the developmental pathway, non-embryonic myogenesis shares a similar molecular and anatomical setup as embryonic myogenesis, but implements co-option and loss of molecular modules.

## BACKGROUND

Musculature is a tissue specialized in contraction shared among all eumetazoans. Its cellular components contain molecular structures based on actomyosin and an array of accessory proteins which allows contractility^1^. Myogenesis operates through a progressive activation of transcription factors organized in a hierarchical and modular network that drive cell fate specification, and is followed by specific cell behaviors that lead to muscle differentiation and organization^2^. For example, trunk skeletal muscles of vertebrates develop from somites and are determined by the expression of the paired box transcription factor Pax3^3^, whereas cardiac, pharyngeal^4^, and possibly also esophagus striated muscles^5^ originate from cardio-pharyngeal mesoderm and are determined by T-Box1. A common origin of heart and pharyngeal muscles from the cardio-pharyngeal field predates vertebrate evolution^4, 6–8^ and has been hypothesized to be a chordate synapomorphy^9,10^. In metazoans, muscles can develop during embryogenesis and during non-embryonic development of adult organisms, i.e. regeneration and asexual development. The myogenic process may be triggered by populations of multi- or unipotent stem cells^11–14^. Post-embryonic myogenesis also occurs in species with indirect development and complex life cycles^15^. These animals undergo drastic changes between their larval and adult bauplan, such as during metamorphosis, and the musculature can radically change architecture within the same organism between different life stages^16,17^.

With their bi-phasic life history and asexually reproducing colonial species, ascidians (Tunicata) offer great opportunities to study the development of different muscle architectures in different life stages^18^. From the fertilized egg, a stereotyped development and largely determinative embryogenesis^19^ leads to the formation of a planktonic larval body. In the larval tail, bands of mono-nucleated myocytes are arranged in a striated fashion^20^ and express a myosin heavy chain, specific to embryonic muscles^21,22^. When the larva settles on a substrate and metamorphoses into a sessile filter-feeding adult, the sarcomeric arranged musculature gets resorbed along with the tail^23^ and non-striated circular and longitudinal body muscles form along the mantle of the organism, together with the cardiac muscles^20,24^. The adult body wall muscle are described as smooth (non-striated) muscles, which apparently evolved due to a loss of sarcomeric organization, probably in order to cope with a sessile lifestyle, which requires slow contractions for the fine tuning of water inflow^20^. The body and heart muscles express two specific, post-metamorphic myosins: myosin heavy chain 3 and myosin heavy chain 2^22^, respectively.

During embryonic development of solitary ascidians, maternal deposition of the zinc finger family member Zic-r.a (Macho-1) is essential for early muscle specification^25^. Zic-r.a activates the expression of the T-box transcription factor *Tbx6*, necessary to induce the tail muscles^26^. Zic-r.a and Tbx6 together with beta-catenin define the cardio-pharyngeal field by activating the bHLH regulatory gene *mesodermal posterior* (*Mesp*)^27^. The cardio-pharyngeal field, a.k.a. the trunk ventral cells (TVCs) in *Ciona,* give rise to both the heart^28^ and part of the adult ascidian body musculature. The expression of the homeobox transcription factor *Nk4*, orthologue to tinman/Nkx2-5, in TVCs antagonizes Tbx1/10 and promotes the cardiac muscle fate through the activation of *Gata4/5/6*^29^. The heart muscle will continue to differentiate and express myosin heavy chain 2^22^. On the other hand, a subset of the TVCs express the transcription factor Tbx1/10, which promotes the expression of *Ebf* (COE, Collier/OLF/Ebf) and orchestrates the transition to the myogenic program by the activation of myogenic regulatory factor(Mrf), eventually giving rise to the longitudinal muscle and the muscles around the atrial siphon^29^. Another set of body muscle, the oral siphon muscles, derive from a different population of cells named trunk lateral cells (TLCs) in *Ciona*. TLCs follow a different fate but are partially regulated by the same transcription factors (TFs) involved in muscle development of TVCs^30,31^. After metamorphosis, all the body muscles continue to differentiate by expressing the same myosin heavy chain 3^22^.

In colonial species of ascidians, such as *Botryllus schlosseri,* the post-metamorphic individual, the oozooid, begins asexual budding from undifferentiated cells, in a process named blastogenesis, which leads to the development of an adult zooid, the blastozooid (Fig. 1). Therefore, the life cycle of a colonial ascidian is characterized by three different body plans: the larva displaying most chordate features, the post-metamorphic oozooid and the asexually propagated blastozooid. By producing multiple buds, in a cyclic manner, blastogenesis eventually leads to the formation of colonies composed of several genetically identical blastozooids. Regardless of their different ontogenetic origin, the overall anatomy of the oozooid and the blastozooid are similar, including the general organization of their musculature^20,24,32^. However, in blastogenesis the process of development is direct, namely it skips the determinative steps of embryonic development and passes neither through a larval stage nor metamorphosis ^33^.

**Fig. 1.**
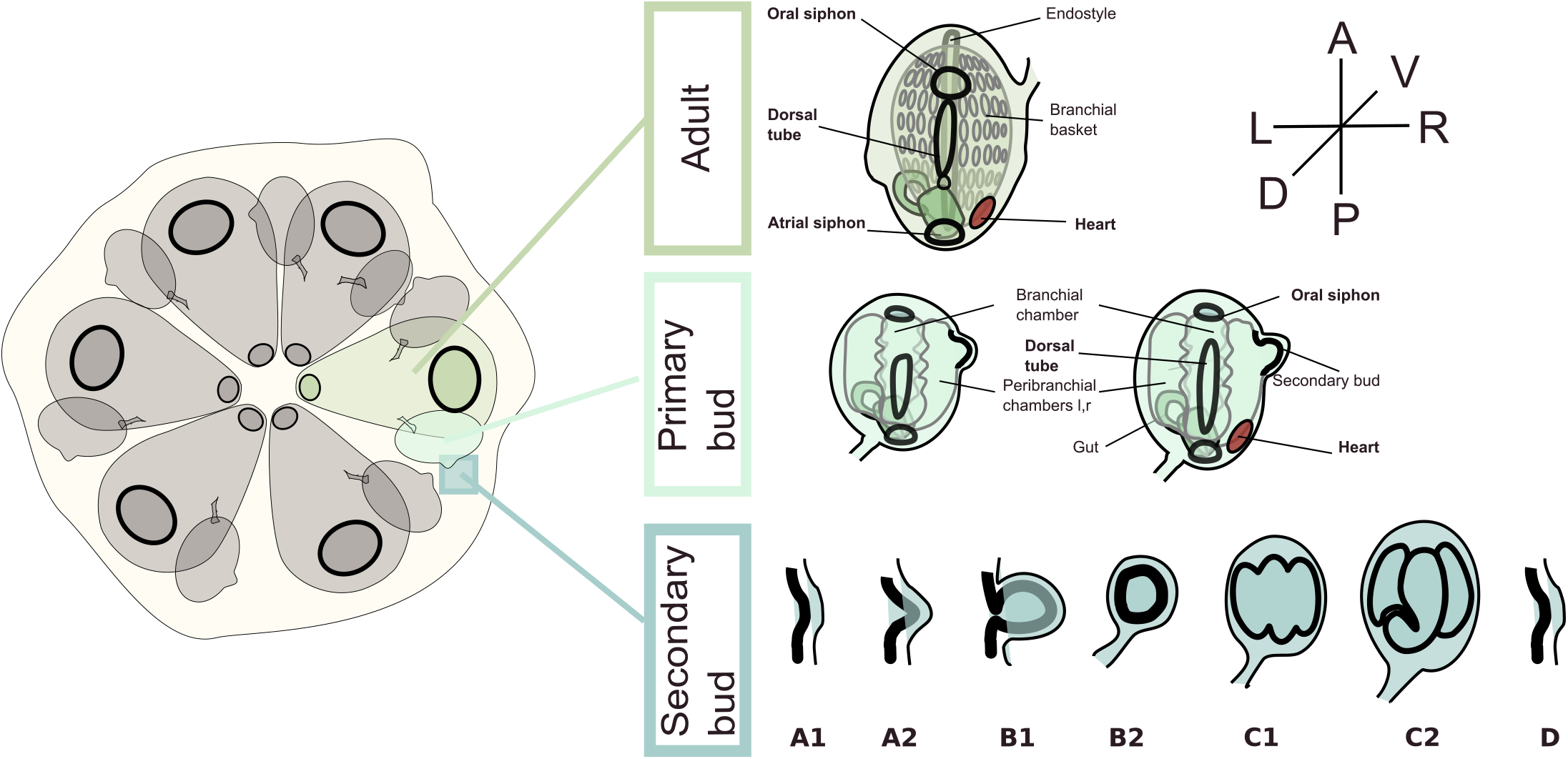
Development and staging of a Botryllus schlosseri colony. The staging of the animals was performed after Lauzon^67^. The secondary bud develops as thickening of the peribranchial epithelium and the epidermis (stages A1, A2), which evaginates and closes forming a double monolayered vesicle (stages B1, B2). The inner vesicle undergoes morphogenesis and is subdivided into three chambers (stages C1, C2). During “takeover” (stages), the adult degenerate and get resorbed, the primary bud become adult, the secondary bud becomes the primary bud and a new blastogenetic cycle begins for the next secondary bud.

In order to investigate the mechanisms underlying clonal replication of muscular systems in colonial ascidians and infer the origin of muscle precursors during blastogenesis, we selected genes involved in embryonic-and-metamorphic myogenesis in the well-studied solitary ascidian *Ciona*, and followed their expression during blastogenesis of *B. schlosseri*. We took advantage of the extensive literature on myogenesis in solitary ascidians, as well as of the broad and detailed descriptions of *Botryllus* ontogeny^34^, to reconstruct the dynamics of muscle formation in ascidians. Our study revealed that during myogenesis of an asexually derived zooid the embryonic and post-metamorphic myogenic genes are only partially co-opted, reflecting a lack of maternal signals and the absence of the larval stage.

## RESULTS

### Changes in muscle architecture during metamorphosis and blastogenesis

To understand whether the molecular structure of the differentiated muscles in colonial ascidians reflects the expression scenario described in solitary species^22^, and in order to compare the molecular structure of the muscle fibers between oozooid and blastozooid, we searched for members of the muscle type class two myosin heavy chains (MYH) in the transcriptomes of multiple stages in *Botryllus schlosseri*^35,36^. Four paralogues were identified in *Botryllus schlosseri*, including three muscle specific (MYH1, MYH2 and MYH3) and one non-muscle specific (MYH 9/10/11/14). The three muscle MYH clustered together with their corresponding homologous sequences of other solitary ascidians belonging to the suborder Phlebobranchia or Stolidobranchia (Supp. Fig.1). MYH3 subfamily constituted the sister group to the cluster of ascidian MYH1, MYH2, and their corresponding vertebrate paralogues. Within the ascidian paralogues, MYH1 and MYH2 clustered as the sister-group and were more closely related to the vertebrate muscle specific MYH sequences (Supp. Fig.1).

*In situ* hybridization revealed that the three mRNAs coding for the muscle relevant MYH proteins are expressed in different regions and at different time points of *B. schlosseri* development. Embryonic *Myh* (*Myh1*) was exclusively expressed in the larval stage in tail muscles (Fig. 2.A). The caudal musculature of the swimming larva consists of cylindrical mono-nucleated cells in longitudinal rows flanking the notochord. During resorption of the larval tail, in metamorphosis, muscle cells lose their cohesion and are pushed into the body cavity, where they become surrounded by phagocytes^37^. The process of tail resorption in *Botryllus schlosseri* (Supp. Fig. 2) occurs within 10 minutes approximately. *Myh1* was expressed in the swimming larval tail in six rows of muscle cells (Supp. Fig. 2.A1-A6). Within cells of the larval tail muscle, the RNA expression was oriented along the anteroposterior axis at the level of the nucleus (Supp. Fig. 2. A5).When regression of the tail was almost complete, the RNA expression became restricted to more anterior sites of the tail (Supp. Fig. 2.B1-2,6), and subsequently formed two separate fields, in the posterior larval trunk (Supp. Fig. 2.B3-4,7). Muscle cell connections were lost and cells became roundish, but myofibrillar striation remained (Supp. Fig. 2.B5). Single cells of embryonic musculature were still found within the mantle of the young oozooid (Supp. Fig. 2.C1-4). The muscle architecture of the zooid, both oozooid and blastozooid, heavily differed from the larval tail musculature. The adult body musculature is characterized by the expression of *Myh*3. In *Botryllus,* the first expression of *Myh3* was only observed during metamorphosis at the anterior end of the young oozooid and where the future intersiphonal muscles were set in place (Supp. Fig. 3.A1-5, B.1-3,C). In the early oozooid, single larval muscle cells were still present in the mantle of the animal (Supp. Fig. 2.C3, Supp. Fig. 3.A3-6,C), but they do not express *Myh3*. The late oozooid, which already carries a bud in stage C2, showed clear dorsal expression of *Myh3* around the oral and atrial siphons, the intersiphonal muscle bands and the body wall musculature (Supp. Fig. 3.D1-4,F), but not on the ventral side (Supp. Fig. 3.E1-4). The fully developed post-metamorphic oozooids expressed *Myh3* in their entire body wall, including the dorsal and ventral musculature, highlighting the fibrous structure of adult muscles (Fig. 2.B): i.e. the circular muscles around the oral and atrial siphons, the longitudinal muscles, and the intersiphonal muscle (Fig. 2.F). The blastozooid (zooid developed by budding) expressed *Myh3* from the primary bud at stage A1 until the fully differentiated zooid (Fig. 2.D). Expression of *Myh3* was detected in circular, longitudinal and intersiphonal muscles, mimicking the patterns observed in the oozooid.

**Fig. 2.**
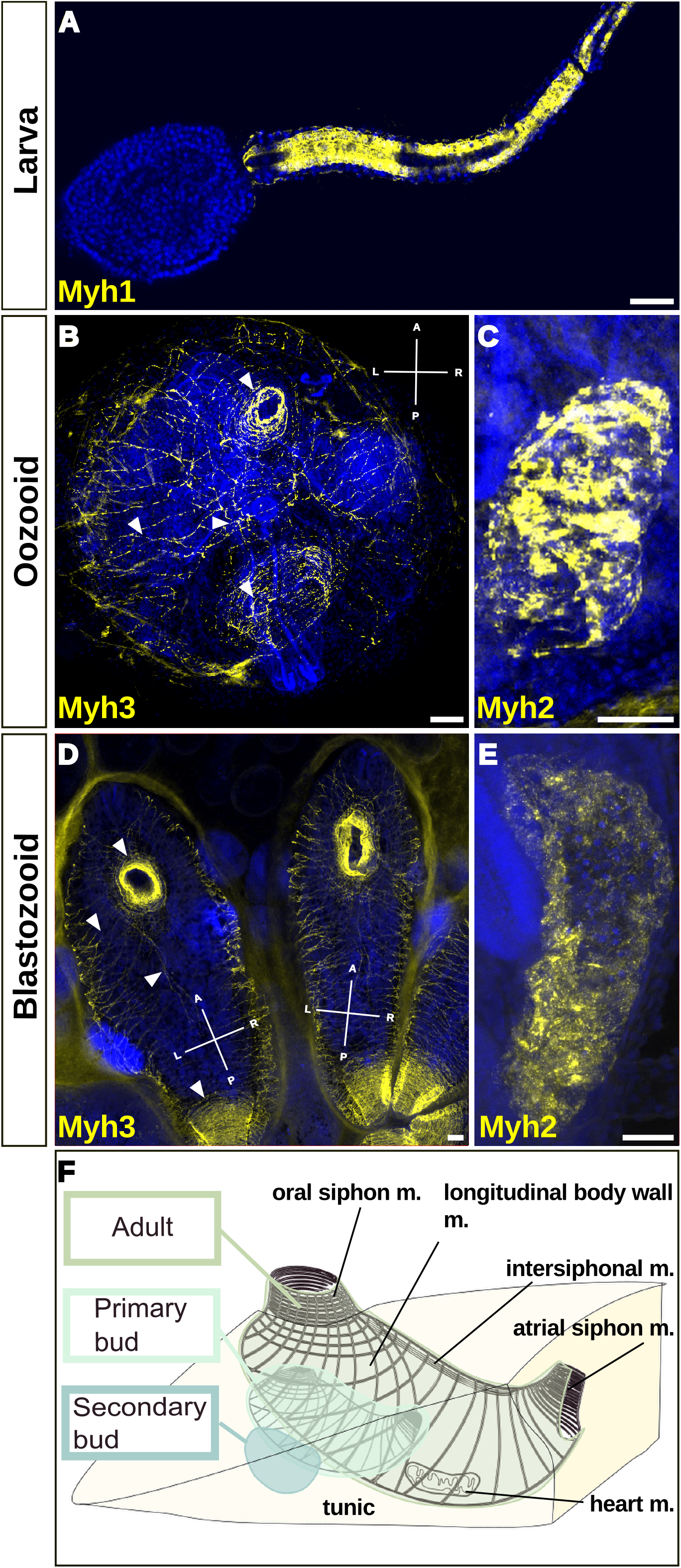
Fluorescent in situ hybridization showing the expression of Myosin heavy chain mRNAs at different stages of Botryllus schlosseri life cycle. (A) Expression of *Myh1* (yellow) along the striated muscle in the larval tail. (B) *Myh3* expression along muscle circular (oral and atrial siphons) and longitudinal muscle fibers in the oozooid body wall. (C) Expression of *Myh2* in the heart of the adult oozooid. (D) *Myh3* expression in the body wall of a blastozooid: circular and longitudinal fibers. (E) *Myh2* expression in the heart of an adult blastozooid. Gene names are indicated in the lower left corner of the pictures in yellow. Arrowheads point circular and longitudinal fibers. Blue nuclei: Hoechst staining. Scale bar: 50 μm. (F) Schematic drawing of the body musculature of a colony in stage C1. Adult, a primary and a secondary bud are embedded in common tunic At stage C1 adult and primary bud the muscle fibers are developed and both are able to contract. The body wall musculature consist of circular muscles around the oral and atrial siphon, longitudinal muscles and a band of intersiphonal muscles, connecting the two siphons. The heart lies to the ventral and right side of the zooid body.

The heart in ascidians consists of a simple two-layered tube (myo- and pericard) that beats haemolymph in a reversible orientation through an open circulatory system^28,38^ and is characterized by the expression of *Myh2*. The heart of solitary ascidians does not fully differentiate or begin contractions before metamorphosis, whereas the heart of some colonial ascidians may become fully functional before settlement^39^. In *Botryllus*, *Myh2* expression was first observed at the ventral side of the swimming larva (Supp. Fig. 4.A1-5). The field displays already a sac-like form. In the early oozooid, *Myh2* expression was observed on the beating heart at the left side of the body (Supp. Fig. 4.B1-5). In the fully developed oozooid heart, the muscle fibers of the heart, expressed *Myh2* (Fig. 2.C). Comparably, *Myh2* expression in blastozooids was localized to the heart of primary buds, and expression was maintained throughout development and in the fully differentiated zooid (Fig. 2.E).

### Partial re-deployment of embryonic myogenic motifs during blastogenesis

To analyze if myogenic motifs expressed during embryonic development and metamorphoses have been co-opted during blastogenesis, we focused on the expression of 17 candidate genes well characterized during the myogenesis of solitary species^31,40^. We first described the embryonic development of the *B. schlosseri* to assay the conservancy of cleavage patterns between solitary and colonial species. We observed that the early cleavage patterns are comparable to the stereotypical patterns observed in solitary ascidians, including the extensively studied *C. robusta*^33,41,42^. However, the timing between each cleavage is longer, and the time from the first cleavage and the larval hatching is around 5 days (Supp. Fig. 5). The expression of *Zic-r.a*, *Tbx6, Ebf* and *Tbx1/10* also confirmed the temporal and spatial pattern of expression observed in solitary species: the mRNA coding for the zinc finger transcription factor Zic-r.a was localized vegetally at the tip of the two blastomeres of the two-cell stage embryo (Supp. Fig. 6.A). At a 110-cell stage the expression localized in two posterior blastomeres (Supp. Fig. 6.B). *Tbx6* mRNAs was expressed in a bilateral fashion (Supp. Fig. 6.C-D) in the presumptive myoplasm and future larval tail muscles (Supp. Fig. 6.E). Bilateral *Ebf* expression has been observed in a group of four single cells at the anterior trunk in an early tail bud stage (Supp. Fig. 6.F).

Next, we carried an initial assessment for presence and relative abundance of gene expression using the transcriptomes obtained from non-fertile colonies at seven blastogenetic stages^43^. These analyses showed the presence of only a subset of candidate genes, whereas others genes important for muscle development, typically expressed early in the embryonic development of *Ciona* were not found to be expressed during blastogenesis (Fig. 3). The absence of expression of early myogenic genes was also supported by RT-PCR (Supp. Fig. 6.G) and FISH (data not shown). The transcripts that were found in a negligibly low copy number or absent from the transcriptomes belong to myogenic transcription factors *ZicL, LIM (Lhx3), Tbx6, Hand-r* and *Mesp,* whereas *Nk4, Tbx1/10, FoxF, Islet, Myh2*, Gata4/5/6, Mrf*, Myh1* and Zic-r.a (*Macho-1*) were expressed at low levels. In contrast, *Myh3, Ebf* and *Ets* were myogenic factors expressed at high levels throughout the seven stages of *Botryllus* blastogenesis.

**Fig. 3.**
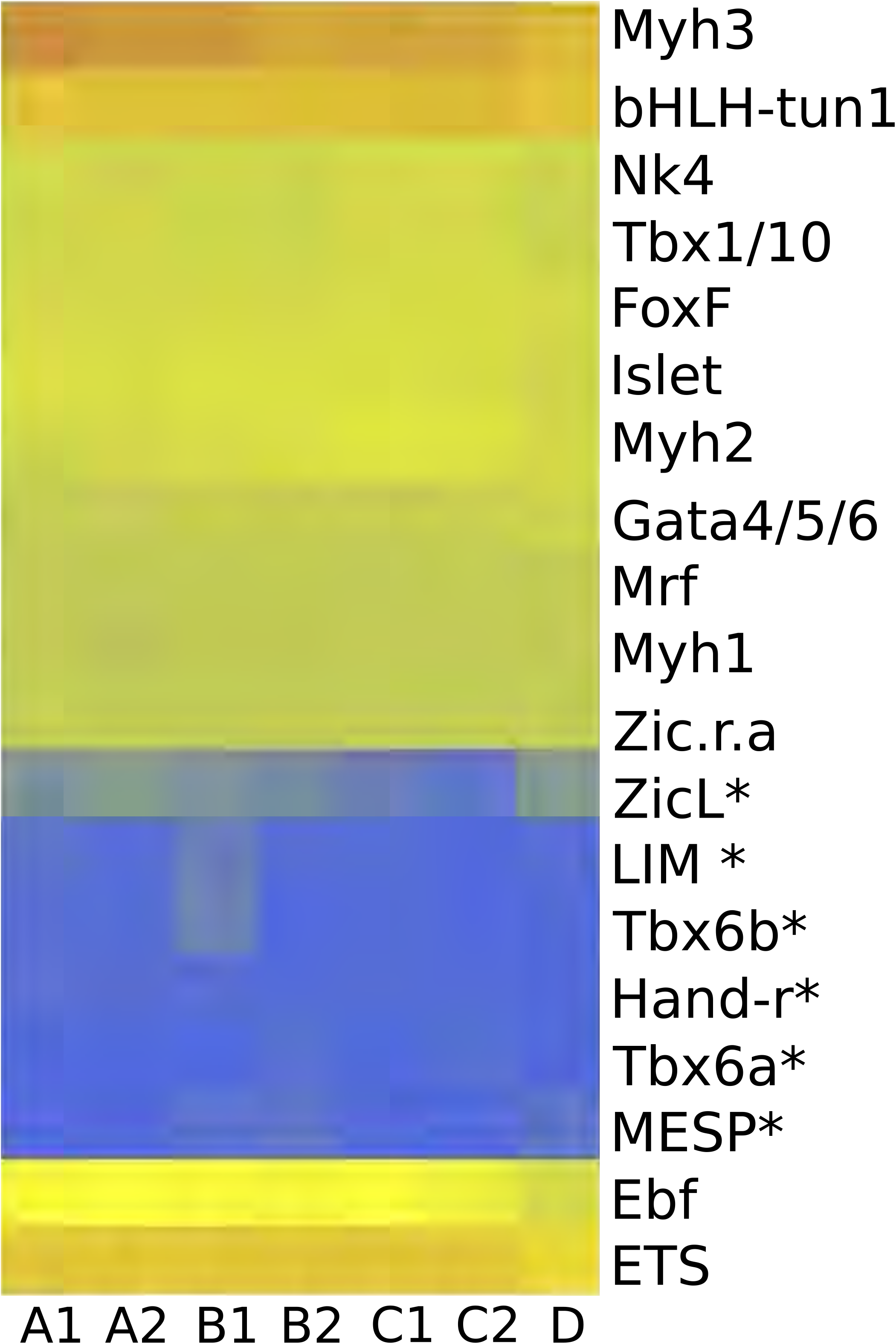
Presence of components of myogenic motif in transcriptomes of blastogenetic stages. Differential mRNA expression level of candidate genes for myogenesis along the seven successive stages of blastogenetic development [A1-D]. Royal blue: RPM<1 (e.g. *Hand-r*), yellow: RPM 1>50 (eg.*Myh2*), coral: RPM >50 (eg. *Ets*), asterisk: not expressed.

### Development of the body wall musculature during *Botryllus* blastogenesis

In order to understand where the body wall muscles originate and develop during *Botryllus* blastogenesis, we investigated the expression of *Tbx1/10*, *Ebf*, and *Mrf*. These three TFs are expressed in one of the two muscle precursor fields in the *Ciona* larval head, specifically in the trunk ventral cells (TVCs). In contrast to descriptions during ascidian embryogenesis, we observed three main regions of expression of these genes in *Botryllus* blastogenesis: (1) a mesenchymal region between the dorsal tube and the dorsal epidermis, (2) the dorsal side of the branchial chamber where the future intersiphonal muscle will develop, and (3) the mantle or body wall, which corresponds to a membrane beneath the tunic made of epithelial tissue, connective tissue, musculature, blood vessels, and nerves (Fig. 4.C,F,I).

**Fig. 4.**
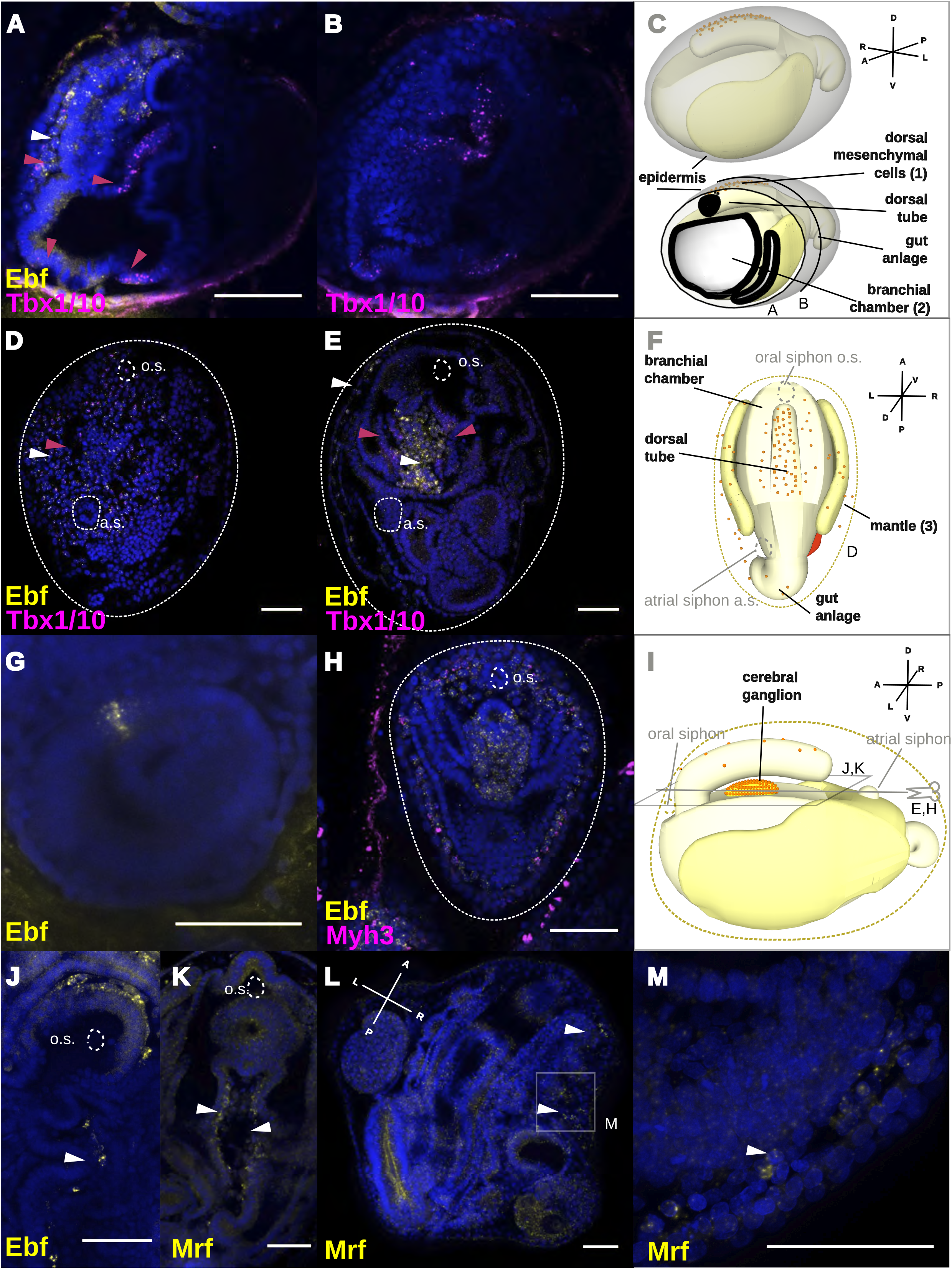
Fluorescent in situ hybridization (FISH) showing expression of body muscle myogenic TFs during blastogenetic development of Botryllus schlosseri. (A-E) Double ISH showing the expression of *Tbx1/10* (magenta) and *Ebf* (yellow). (A-B) *FISH* preformed on the same animal in stage C2-D. (A) Magenta arrowheads point to the expression of *Tbx1/10* in the dorsal mesenchyme, the dorsal region of the branchial basket and the ventral region of the branchial basket lateral to the endostyle, white arrowheads point to the expression of *Ebf* in the dorsal mesenchyme. (B) Posterior and transversal section of the same sample showing the expression of *Tbx1/10* in the dorsal and ventral branchial basket. (C) Schematic representation of a developing Botryllus secondary bud in stage C2-D, showing the dorsal tube the branchial chamber, the gut and the epidermis. (D-E) FISH preformed in stage A1. (D) Magenta arrowheads point to the expression of *Tbx1/10*, white arrowheads to the expression of *Ebf* in the mantle. O.s: oral siphon, a.s.: atrial siphon. (E) Magenta arrowheads point to the bilateral expression of *Tbx1/10* in the intersiphonal region, white arrowheads to the expression of *Ebf* in the mantle and the forming cerebral ganglion. (F) Schematic representation of a developing *Botryllus* primary bud in stage A1. (G) Earliest expression of *Ebf* in few single cells of the vesicle like secondary bud in stage B2, oriented to the primary bud and hence the future dorsal side. (H) Double ISH showing the expression of *Ebf* (yellow) and *Myh3* (magenta), *Ebf* is expressed in mantle together with *Myh3*, in the cerebral ganglion and in the dorsal tube. (I) Schematic representation of a developing *Botryllus* primary bud in stage A, sagittal view. (J) *Ebf* expression in the region of the intersiphonal muscle. (K) Max projection showing the expression of *Mrf* in the region of the intersiphonal muscle. (L-M) *Mrf* expression in the mantle of the primary bud. Blue nuclei: Hoechst staining, dashed line: outline of primary bud, scale bar: 50 μm. Asterisk: unspecific staining in either tunic, gonads, or insufficiently bleached auto-fluorescent cells.

*Tbx1/10* expression began in the secondary bud at stage C2, in mesenchymal cells between the dorsal tube and the epidermis (Fig. 4.A). In the early primary bud (between stage D-A1), *Tbx1/10* was expressed in mesenchymal cells distributed underneath the epidermis along the entire body of the zooid, i.e. the mantle (Fig. 4.D). At stage C2 the secondary bud started to express *Tbx1/10* ventrally in the branchial chamber flanking the forming endostyle as well as in its medio-dorsal region (Fig. 4.A-B). This second domain of expression appeared to concentrate laterally in the epithelium and into the cerebral ganglion region in stage D/A1 of the primary bud (Fig. 4.E), and was maintained until A2, as it began to appear more diffuse. *Tbx1/10* was also expressed on the ventral side of the bud within the region of the forming heart (see below).

The earliest expression of *Ebf* was in a domain composed of 3 to 5 cells in the secondary bud at stage B2 (Fig. 4.G). A second domain of expression appeared in the secondary bud on the dorsal tube^34^ at stage C1 and in mesenchymal cells between the dorsal tube and the overlying epidermis, where it was co-expressed with *Tbx1/10* (Fig. 4.A). In the primary bud at stage D-A1, *Ebf* was co-expressed with *Tbx1/10* in mesenchymal cells along the mantle of the zooid (Fig. 4.D). From stage B1/B2, *Ebf* was expressed ubiquitously in the cerebral ganglion, in scattered cells around the forming neural gland, and it was co-expressed with the body wall muscle marker *Myh3* in the mesenchymal cells along the mantle, where the future muscle fibers formed (Fig. 4.H). In a C2 primary bud, *Ebf* was expressed in few scattered cells along the intersiphonal region (Fig. 4.J). The transcriptomic data we analyzed showed a low expression of *Mrf* in all blastogenetic stages (Fig. 3.H). In addition, FISH showed feeble expression of *Mrf* only in primary buds at stage A2, in cells of the mantle (Fig. 4.L-M), as well as in a dorsal domain of the branchial chamber, possibly where *Tbx1/10* was also expressed (Fig. 4.K). However, due to the low expression of *Mrf* RNA, it was not possible to perform double FISH for *Mrf* and *Tbx1/10* simultaneously.

### Presence of putative muscle stem cells in the zooid

In *Ciona,* the atrial and oral siphon muscles have been shown to maintain a population of putative muscle stem cells that is Mrf-/bHLH-tun+^31,44^. While *Mrf* is expressed in differentiating muscle cells, the tunicate-specific helix-loop-helix transcription factor *bHLH-tun* is expressed in muscle stem cells ^31,44^. In the *Botryllus* oozooid, the *bHLH-tun* mRNA was expressed in a ring around the two siphons (Supp. Fig. 7.A1-5, B.1-4), which probably corresponds to the inner muscle population of the siphon, as was previously suggested in *Ciona*. In addition, *bHLH-tun* was expressed in a few cells of the lateral body wall (Supp. Fig. 7.A2) and in a few cells on top of the dorsal tube (Supp. Fig. 7.C1-4). In the *Botryllus* blastozooid, such spatial pattern of expression was not been detected, however transcriptome analyses showed that *bHLH-tun* is in fact present during different stages of blastogenesis (Fig. 3).

### Heart development during blastogenesis

In order to investigate the formation of heart musculature during blastogenesis we followed the expression of the TFs *FoxF, Tbx1/10* and *Nk4*, which characterize the TVC embryonic lineage that gives rise to the heart muscles in *Ciona* ^29,40^ (Fig. 5). In contrast, *Botryllus FoxF* mRNA was expressed in cells of the bud epidermis throughout blastogenesis (Fig. 5.A-C), and more specifically on the dorsal side of the branchial chamber of the secondary bud on stage D (Fig. 5.A). In the *Botryllus* primary bud, *FoxF* seemed to concentrate at sites lateral to the cerebral ganglion (data not shown). Last, we detected expression of *FoxF* in a cluster of cells in the left ventral side of the mantle where the heart formed (Fig. 5.*B*), and in the heart of primary buds (Fig. 5.C). *Tbx1/10* transcripts showed similar patterns of expression to those described above for *FoxF*, but in addition *Tbx1/10* showed expression on the secondary bud (stage D/A1) in a cluster of cells on the left ventral side of the mantle where the heart presumably will form (Fig. 5.D)., Heart field determination in *Ciona* is mainly characterized by the expression of *Nk4*, which antagonizes *Tbx1/10* and *Ebf*^29^. In the *Botryllus* secondary bud at stage C2, *Nk4* was expressed asymmetrically on the left side of the branchial chamber epithelium and in the entire left peribranchial chamber (Fig. 5.E). At stage B1, *Nk4* expression was observed in the forming heart (Fig. 5.F). Expression of *Nk4* in the myocardium remained high during the complete development of the heart until the myocardium was separated from the pericardium (Fig. 5I). To test whether *Ebf* and *Nk4* expression domains are mutually exclusive, we performed double ISH (Fig. 5.G-H) on a single colony. *Nk4* was expressed all over the secondary bud at stage B2-C1 except for the region of *Ebf* expression, which corresponds to the site where the dorsal tube will form (Fig. 5.G). We never found *Ebf* expressed at any stage of ascidian heart development (Fig. 5.H).

**Fig. 5.**
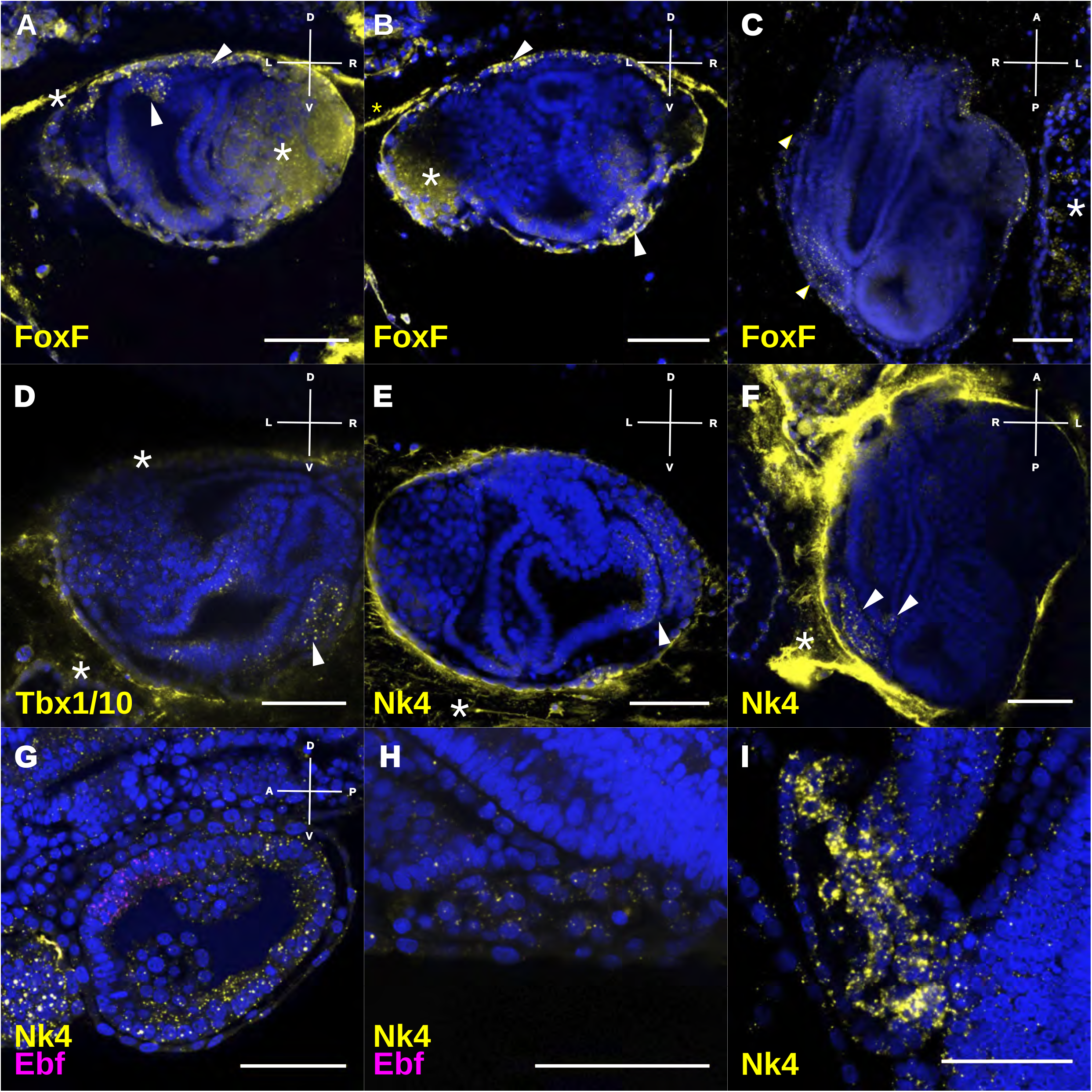
Expression of TFs heart development-related revealed by FISH. (A-C) Expression of *FoxF* in secondary bud at stage D: (A) in the dorsal branchial chamber and epidermis, (B) in the forming heart vesicle and epidermis and (C) *FoxF* expression in the heart and epidermis in a ventral view shown by a max projection of 13 slides, stage A2. (D) *Tbx1/10* expression in stage D heart. (E) Asymmetric *Nk4* expression in stage C2, over the entire right peribranchial chamber (white arrowhead) and the right side of the branchial chamber. (F) *Nk4* expression in a ventral view of a primary bud in stage B1. (G-H) Double FISH with *Nk4* and *Ebf* in the same B2-C1 colony (G) in the secondary bud and (H) the heart of the primary bud. (I) *Nk4* restricting to the myocardium in late primary bud. Probes are indicated in the lower left corner of the pictures in the same color of the expression. Blue nuclei: Hoechst staining, dashed line: outline of primary bud, scale bar: 50 μm. Asterisk: unspecific staining in either tunic, gonads, or insufficiently bleached auto-fluorescent cells.

## DISCUSSION

In this study, we characterized mRNA expression patterns of myogenesis-related genes during the two different ontogeneses of the colonial ascidian *Botryllus schlosseri*. By following the sequential expression patterns of myogenic TFs throughout a complete series of developmental stages, we reconstructed the putative cellular precursors for body and heart muscles. We show that, within a single chordate species, the myogenic transcriptional motifs are only partially co-opted and cellular origin and transcriptional regulation that lead to adult muscles are coordinated differently during embryonic and non-embryonic developmental processes.

### Colonial and solitary embryogenesis/metamorphosis share myogenic motifs

Comparative analyses of expression patterns of myogenic TFs during embryogenesis showed that several genes are conserved between *Botryllus* and the solitary species studied so far^45–48^ suggesting that core elements of the myogenic regulatory cascade of solitary species are conserved during the sexual development of colonial species. For instance, the maternal expression of the posterior determinant and muscle specifier *Zic-r.a* (*Macho-1*) ^45,49^ in *B. schlosseri,* as well as its target *Tbx6*, indicate that the upstream determinants governing ascidian muscle development share the same expression as that reported for solitary forms. *Ebf was* also expressed during embryogenesis in *B. schlosseri*, and *Mesp* showed some expression despite the fact that it was not detectable by ISH, which we attribute to a temporal restriction of *Mesp*. Our data support the observation of Ricci et al. (2016)^33^ showing that the early cleavage pattern seems to be conserved between solitary and colonial ascidians, at least for *B. schlosseri*. These data confirm the robustness of the ascidian expression and cleavage patterns across solitary and colonial species despite large variation in egg size, and strongly suggests that features of developmental mechanisms responsible of larval and post-metamorphic myogenesis might be well conserved in the whole class.

### Differentiated muscle cells share molecular components in oozooids and blastozooids

In *Botryllus*, the body and heart muscles are formed anew, starting from the post-metamorphic oozooid and in every adult blastozooids during each blastogenetic cycle^20,24^. Regardless their different cellular origin and their divergent ontogenies, the oozooid and the blastozooid of *Botryllus schlosseri* present a similar arrangement of body structures, tissues, and cell types^32^. A few differences among the two zooid types, include differences in body size (the oozooid generally being bigger than the blastozooid), differences related to the architecture of the branchial basket^34,50^, and the organization of the musculature, e.g. the number of muscle fibers varies as well as their arrangement in the atrial siphon^24^. Despite the observed variation in muscle organization, both oozooid and blastozooid muscle cells expressed the same myosins in their fully differentiated fibers, i.e. *Myh3* in the entire body wall musculature, and *Myh2* in the heart. These results confirm that muscles of solitary and colonial ascidians express the same genes during final differentiation of muscle cells. Furthermore, within distinct zooids of the same colonial species, different developmental trajectories lead to a similar differentiation process and similar types of muscles.

### Regulatory factors of myogenesis are only partially co-opted during blastogenesis

The transcriptomic profiles of the blastozooid developmental stages showed the presence of only a subset of the myogenic TFs engaged during ascidian embryogenesis. With the exception of *Zic-r.a*, upstream regulators of embryonic myogenesis, such as *Tbx6*, *LIM (Lhx3), ZicL, Mesp*, and *Hand-r* are not expressed in blastogenesis. During *Botryllus* embryogenesis, Zic-r.a is expressed in the neural plate, and during blastogenesis, it is expressed in the dorsal tube (Supp. Fig. 8). *Zic-r.a* expression in blastogenesis suggests a neurogenic rather than a myogenic function of *Zic-r.a* (Prünster et al, *submitted*). For instance, its expression does not seem linked to *Tbx6*, one of its downstream myogenic targets, which is absent in blastogenesis. This neurogenic role has been described in *Ciona* where, beside its function in early myogenesis, *Zic-r.a* is also zygotically expressed during neurogenesis, as suppressor of notochord fate ^51^. The direct development of the bud, which lacks a larval stage with a notochord and tail musculature, may explain the absence of the early muscle transcriptional module. However, a lack of myogenic upstream regulators did not prevent the expression of late myogenic TFs, which are co-opted during blastogenesis. Precisely the same way that TFs are expressed in the TVCs of the *Ciona* larva throughout cardio-pharyngeal development^40^. These results suggests a degree of plasticity in the regulation of myogenic transcriptional modules in ascidians, which can be decoupled from the control of maternal determinants and early zygotic transcriptional regulators.

### De novo origin of musculature in blastogenesis

In blastogenesis, the body muscle fate seems to be regulated by a kernel of genes that are expressed in the dorsal domain of the developing bud. In particular, *Tbx1/10+* and *Ebf+* cells are localized in a set of mesenchymal cells between the dorsal tube and the epidermis. These cells have been previously described in *Botryllus* and *Diplosoma* as neural precursors, which migrate and cluster forming the cerebral ganglion^52,53^. However, the sequential expression -both temporal and spatial- of *Tbx1/10*, *Ebf* and *Mrf* during bud development suggests that these cells migrate from the dorsal tube towards the lateral mantle, aligning where the future muscle fibers form There precursors might end up in the circular musculature of both siphons, as well as in the longitudinal muscle of the mantle. Therefore, in addition to center of neurogenesis, the dorsal tube could also have an additional role in myogenesis. In *Ciona* the oral siphon muscles do not derive from TVCs but from a population of TLCs, where *Ebf* is first expressed regulating downstream expression of *Tbx1/10* ^30,31^. Without proper functional tests to dissect the interactions between *Ebf* and *Tbx1/10,* we cannot conclude whether the oral siphon musculature network of solitary ascidians is retained during blastogenesis. The presence of muscle stem cells cannot be completely ruled out due to the lack of detection of Mrf-/bHLH-tun+ cells by FISH in the blastozooids, because the colony transcriptomes revealed an expression of bHLH-tun. It remains to be illuminated whether a short-term, renewed every blastogenic cycle, or a long-lived population of muscle precursors persists, which simply remains undetectable by *in situ* hybridization so far.

Another domain of expression of *Tbx1/10-Ebf-Mrf* is located in a dorsal part of the branchial basket epithelium. In this domain, the temporal sequence of gene expression is different. *Eb*f is detected in the overhead mesenchymal cells during later developmental stages, and surprisingly *Mrf* is expressed before *Ebf*. The deviating patterns of expression of these genes as a result of heterochrony are suggestive of cell independent behaviors that occur at the site where the intersiphonal muscles will form. The intersiphonal muscles are the last muscles to form in the *Botryllus* blastozooid and connect longitudinally to the two siphons^24^. The expression of *Tbx1/10, Ebf* and *Mrf* suggests a putative myogenic source in the dorsal epithelial domain, likely linked to the specification of intersiphonal muscles. However, without live tracking, the dynamic nature of muscle formation cannot be fully understood, and we cannot rule out that the *Mrf*+ cells originate from other domains. Because muscle cell fates from these precursors have not been reported in *Ciona*, nor could we detect any such muscle fates in other closely related styelid species^54^ (Supp. Fig. 9), the evidence so far suggests a single origin of myogenic precursor cells in the dorsal domain of buds as a synapomorphy of the Botryllidae.

### During blastogenesis heart muscle origin is possibly uncoupled from body muscles

During blastogenesis, no evidence showed any morphological recapitulation of embryonic heart development, i.e. no ventral fusion of bilaterally located heart progenitors^28^. Instead, the heart either originate from mesenchymal precursors that cluster in the ventro-lateral side of the forming zooid^20,55^ or arise in the same area from the evagination of the branchial chamber^56^. The heart lineage markers, *Foxf, Nk4, Gata4/5/6*^33^ as well as TFs *Tbx1/10* are expressed in the ventro-lateral portion of the branchial chamber, where the cardiac muscle forms. Particularly the late *Nk4* expression is restricted exclusively to the myocardium. While it remains difficult to live-track heart precursor cells and to find their origin, a correlation between the hierarchy of TFs expression and blastozooid organogenesis suggests that the heart is specified “in situ” in the ventro-lateral heart domain. The early expression of *Tbx1/10* together with *Nk4,* that leads to *Nk4* and *Gata4/5/6* expression in the heart primordium resembles the formation of the second heart field^40^. However, the lack of functional relationships between TFs and the dramatically different ontogenesis suggests again a potential re-shuffling of the embryonic cardiac module.

## CONCLUSIONS

In summary, the correlation between patterns of expression and morphogenesis presented here suggests the presence of three putative myogenic domains during *Botryllus* blastogenesis (Fig. 6.A): 1) muscle precursors delaminate from the dorsal tube and migrate along the mantle to form the circular muscles of the siphons and body wall muscles; 2) intersiphonal muscles originate from a dorsal portion of the branchial chamber and seem to be regulated in a different way from the other body wall muscles; and 3) the heart is formed from another population of founder cells localized in a ventro-lateral domain of the bud, which remains poorly characterized. The origin of muscle cells in B. schlosseri blastogenesis is elusive but specifier of cardiac and adult body wall muscles appear partially coopted (Fig 6. B).

**Fig. 6.**
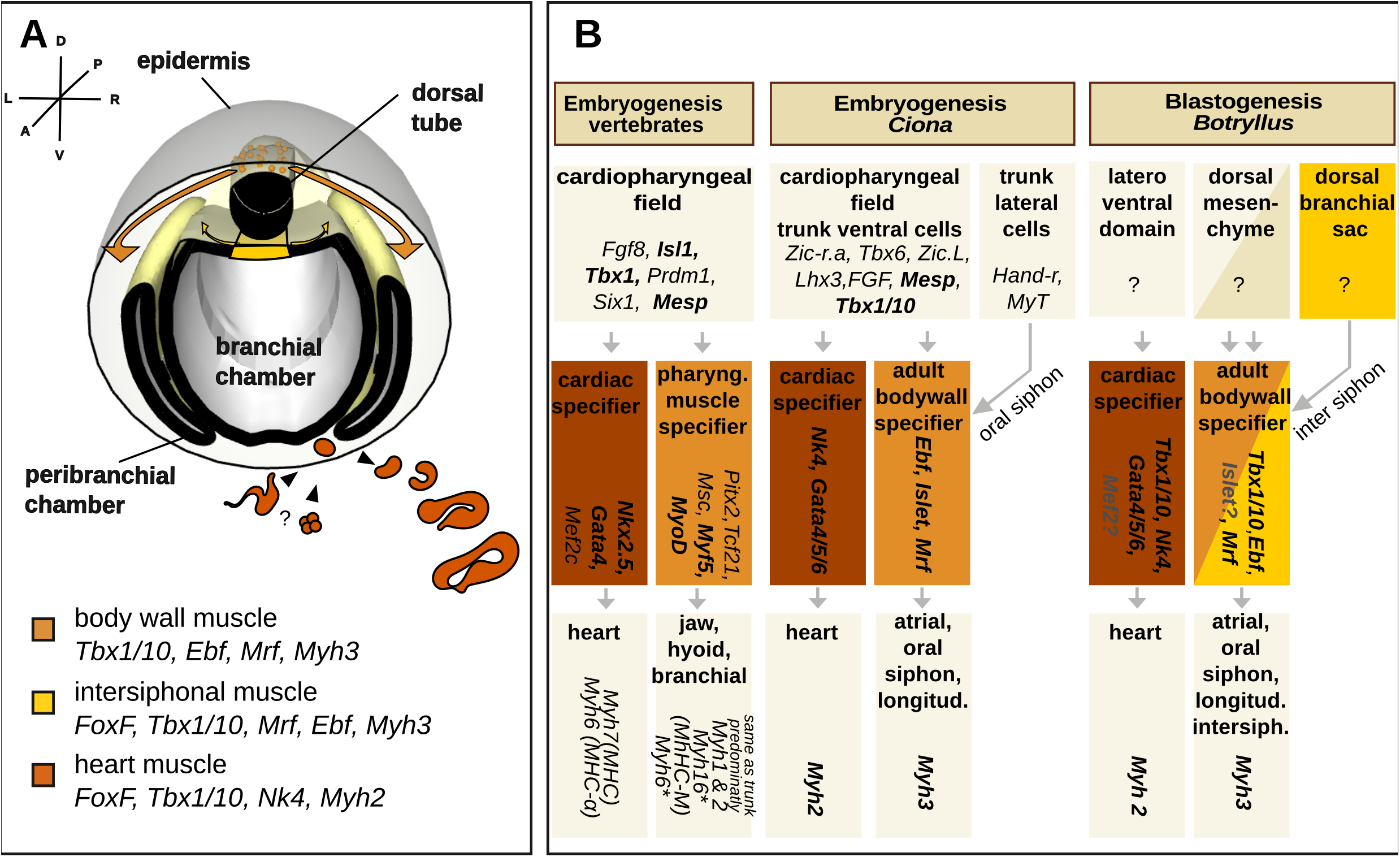
Proposed model showing a modular nature of muscle development. (A) Proposed model for *Botryllus schlosseri* muscle development. Three myogenic regions are depicted: the body wall muscles derived from mesenchymal cells that evaginated from the dorsal tube, the intersiphonal muscles from the posterior branchial chamber and the heart formed by either evagination or clustering of mesenchymal cells in the ventrolateral region. The myocardium formed by invagination, hence the two layered tubular heart of the adult is put in place. (B) Comparison of the cardio-pharyngeal muscle network in chordate development. The expression of genes has been assayed in multiple vertebrates^3, 68–71^, *Ciona intestinalis*^31,40^ and *Botryllus schlosseri*. A number of similarities characterize the inductive signals specifying the cardio-pharyngeal field in vertebrate and ascidian embryogenesis, whilst in blastogenesis a common origin of body wall muscles and heart muscles is unlikely. The cardiac lineage expresses *Nk2.5* and *Gata4* in the forming vertebrate hearts as well as their orthologues in both ascidian species. For what concern *Mef2* expression no information are currently available in ascidians ^72^. An orthologue to vertebrate *Mef2*, is present in the transcriptomic dataset of blastogenesis. Pharyngeal muscle start differentiation by activating the paralogues *Myf5* and *MyoD*; only one orthologue, *Mrf*, is found in ascidians. To activate such in ascidians *Ebf* is expressed in the body muscle lineage. Two myosin heavy chain forms characterize the vertebrate heart. Pharyngeal muscles do not express the same isoforms in all vertebrates: ruminants and rodents express the same myosins as in trunk and limb, namely *Myh1* & *2*, in other animals *Myh 6* and *Myh16* can be expressed in addition. In ascidians, *Myh2* is expressed in the heart, and *Myh3* in the body wall, their proteins are not orthologues of vertebrate Myh2 and 3 (Supp. Fig. 1.). Bolt text indicates conserved expression within at least two species, gray only transcriptomic data, asterisk only in some species.

## Methods

### Animal husbandry

*Botryllus schlosseri* colonies were raised on a 50×70×1 mm glass slides as described previously^61^. A *Botryllus* colony consists of three coexisting asexual generations: the adult filtering zooids, their buds, called primary buds, and the secondary buds (or budlets), sprouting from the primary buds (Fig. 1). Budding (blastogenesis) was staged according to Lauzon et al. (2002)^62^. First a budlet appears as thickening of the peribranchial chamber and overlying epidermis of the adult zooid (stage A), thus closes off forming a double vesicle connected with the parental zooid by the epidermis (stage B). Then organogenesis begins and the inner vesicle separates into three chambers: one central branchial and two lateral peribranchial chambers (stage C). Staging of the development of the embryo has been done according to Conklin^63^.

### Embryo harvesting and dechorionation

Embryos were harvested at different developmental stages from the colony by opening the Botryllus adults with a syringe. Dechorionation was performed in fertilized eggs by shaking the eggs at 60RPM at room temperature in 0.2 % trypsin and 20 mM TAPS, pH 8.2 in seawater for 1.5-2 hours followed by several seawater washes. To calculate the cleavage time, non-dechorionated fertilized eggs have been harvested and kept in filtered seawater at 17°C.

### Gene identification and phylogenetic analyses

RNA sequences where retrieved by tblastn from the *Botryllus schlosseri* transcriptome database http://octopus.obs-lfr.fr/public/botryllus/blast_botryllus.php, full length sequences of proteins were retrieved from Aniseed (https://www.aniseed.cnrs.fr/), and from NCBI. Alignments have been compiled using MAFFT, and sequence trimmed by the TrimAl Gappyout method. Maximum likelihood trees were compiled using PhyML^64^ (Supp. Fig. 10-13, Supp. Text 1).

### Fluorescent in situ hybridization (FISH)

Primers for antisense mRNA probes were designed in the translated region of each gene (Supp. Table 1) FISH was carried out as previously described in Ricci et al. (2016) with the following modifications: 1% Dextran sulfate was added to the Hybridization buffer and the revelation solution. The anti-Digoxigenin Antibody (HRP) (Roche,11207733910) was pre-adsorbed for 1 hour in hybridization solution with a mix of fixed colonies at different stages. When the tunic was exhibiting a very strong background, the animals were manually removed from x the tunic after rehydration, post fixed in 4% PFA for 1h and transferred into washing baskets in 24-well plates. DIG-probe detection was performed with bench-made FITC-Tyramide by 3h incubation. For double FISH, the hybridization of DIG labeled and Fluorescein labeled probes was performed at the same time, fluorescein probes where detected with Cy3-Tyramide. The ISH on embryos was performed after Christiaen et al. (2009)^65^.

### Transcriptional data analysis

Transcripts of RNA-seq data of seven stages of an unfertile colony of *B. schlosseri* SB802d^43^ were quantified by pseudo alignment via Kallisto^66^ to the mixed stage transcriptome database http://octopus.obs-fr.fr/public/botryllus/blast_botryllus.php. Heat map was generated using R, a cutoff for not expressed genes was chosen <1 RPM, lowly expressed 1-50, and high expressed >50 RPM. Orthologues of muscle determinants have been selected by reciprocal blast with published data from NCBI. For some candidate genes, the orthology has been also assessed by phylogenetic analyses (Supp. Fig. 10-14, Supp. Text 1).

### Imaging

Imaging of the NBT/BCIP ISH have been acquired with Zeiss Axio Imager A2 with a 20x magnification, DIC, and color camera. Confocal images were acquired a Leica SP8 (40x/1.1 Water WD 0.6 HCX PL APO CS2) and processed with ImageJ and Inkscape.

## Acknowledgements

We would like to thank M. Khamla for his contribution with the artwork, S. Lotito and L. Giletta for technical assistance. We thank Dr. A. Alie and M. Scelzo for comments. This work was supported by ANR (ANR-14-CE02-0019-01), FAPESP Jovem Pesquisador grant (JP 2015/50164-5) and CNRS/PICS. L.R. was supported by an FRM grant (#FDT20140931163).

## Author contributions

M.M.P, L.R., F.D.B. and S.T. conceived and designed the study. M.M.P. and L.R. performed the experiments. M.M.P and S.T analyzed the data and wrote the manuscript. All authors read and approved the final version of the manuscript.

## Additional information

The authors declare no competing interests that might be perceived to influence the results and/or discussion reported in this paper.

## Supplementary material

***Supp. Fig. 1*** ML tree of class 2 type Myosin heavy chain

***Supp. Fig. 2*** Myh1 expression in larva and during metamorphoses

***Supp. Fig. 3*** Myh3 expression in *B. schlosseri* oozooid

***Supp. Fig. 4*** Myh2 expression in swimming larva and oozooid

***Supp. Fig. 5*** Embryonic development *Botryllus schlosseri*

***Supp. Fig. 6*** Expression of myogenic factors in early embryo

***Supp. Fig. 7*** Expression of bHLH-tun in oozooid

***Supp. Fig. 8*** Zic-r.a expression in embryo, oozooid and blastozooid

***Supp. Fig. 9*** Phalloidin in *Polyandrocarpa zorritensis* (Styelidae) adult, showing absence of intersiphonal muscle bands

***Supp. Fig. 10*** ML tree of Nk4

***Supp. Fig. 11*** ML tree of MRF

***Supp. Fig. 12*** ML tree of Mesp

***Supp. Fig. 13*** ML tree of Ebf

***Supp. Text 1*** Sequences for Phylogeny

***Supp. Table 1*** Primer sequences

